# Causal mapping of self-motion networks in the human brain

**DOI:** 10.64898/2026.09.19.752147

**Authors:** Zoé Dary, Stanislas Lagarde, Samuel Medina Villalon, Hugo Dary, Jacques Léonard, Fabrice Bartolomei, Christophe Lopez

**Author notes:** Address for correspondence: Dr Christophe Lopez, Centre de Recherche en Psychologie et Neurosciences (CRPN), Centre National de la Recherche Scientifique (CNRS) and Aix Marseille University Centre Saint-Charles, Fédération de Recherche 3C – Case B. 3, Place Victor Hugo. 13331 Marseille Cedex 03, France.

## Abstract

Although functional neuroimaging studies using caloric and galvanic vestibular stimulation have identified a distributed cortical network involved in human vestibular processing, including the posterior insula, parietal operculum, temporo-parietal junction, cingulate and frontal areas, the causal contribution of specific brain regions to vestibular self-motion perception remains poorly understood. Invasive electrical brain stimulation (EBS) during stereoelectroencephalography (SEEG) offers a unique opportunity to causally map the cortical networks underlying vestibular self-motion perception in humans.

Here, we retrospectively analyzed EBS-induced vestibular percepts in 354 patients with drug-resistant epilepsy undergoing SEEG. A total of 19,708 stimulations (11,004 at 50 Hz and 8,704 at 1 Hz) yielded 3,015 clinical responses. Vestibular self-motion illusions were defined as sensations of vertigo, dizziness, whole-body rotation, or translation occurring without corresponding physical movement. Stimulation sites were assigned to the seven large-scale functional networks of the Schaefer-Yeo atlas to characterize the network organization of vestibular self-motion perception.

Vestibular self-motion sensations were elicited in 46 patients (21 females; mean age ± SD: 25 ± 11 years) during 86 EBS delivered outside epileptogenic and lesional regions. Percepts ranged from nonspecific vertigo and dizziness (54.7%) to more explicit rotational (30.2%) and translational (15.1%) self-motion illusions. Vestibular responses were most commonly evoked by stimulation of the insula, medial temporal regions, cingulate cortex, inferior frontal gyrus, and premotor cortices. Network-level mapping using the Schaefer-Yeo atlas revealed a non-uniform distribution of vestibular sites across large-scale functional networks, with the highest representation within the salience/ventral attention, visual, and dorsal attention networks.

Together, these findings provide causal evidence that vestibular self-motion perception emerges from activity within a distributed cortical network involved in attentional control and multisensory processing. Beyond advancing our understanding of human vestibular processing, these findings may have clinical relevance for disorders involving altered self-motion perception, including vestibular epilepsy and functional neurological disorders.

## Introduction

The vestibular system is one of the most ancient and important neurosensory systems for encoding and processing self-motion and self-orientation in space. Anatomically, it is highly complex, extending beyond the inner ear’s motion sensors to encompass a vast network of multisensory structures involved in motion and orientation processing^1^. Self-motion and orientation information from these sensors is directly transmitted to the vestibular nuclei in the brainstem and the cerebellum^2^. Multiple parallel pathways then relay self-motion signals to several thalamic nuclei, the basal ganglia, and an extensive cortical vestibular network^3,4^.

Unlike other sensorimotor cortical areas, there does not appear to be a single primary cortical region dedicated to vestibular processing. Instead, vestibular signals are integrated within a highly interconnected cortical network^5^. Single-cell recordings in nonhuman primates have revealed vestibular responses in regions such as the parietoinsular vestibular cortex (PIVC)^6–8^, the ventral and medial intraparietal areas (VIP and MIP)^9,10^, the posterior cingulate cortex^11^, the medial superior temporal area (MST)^12^, and the hippocampus^13^. These regions exhibit varying tuning properties and sensitivities to vestibular and visual self-motion encoding^14,15^ and collectively process self-motion perception according to Bayesian integration principles^16^. Anatomical studies in nonhuman primates, employing tracer injections in the thalamus and cortical regions of this network, have shown that these areas are densely interconnected^5,17^. Notably, the PIVC appears to serve as a hub within this cortical network, as it is more extensively connected to other cortical regions involved in vestibular information processing^5,17^.

The neuroanatomy of the human cortical vestibular network has advanced significantly with non-invasive functional neuroimaging (reviewed in Ref.^18,19^). Stimulation of vestibular sensors or the vestibular nerve using caloric, acoustic, or galvanic vestibular stimulation has been used to map the associated blood-oxygen-level-dependent (BOLD) responses. Consistent with anatomical data from nonhuman primate tracing studies, fMRI and PET approaches have revealed robust responses during vestibular stimulation in a wide range of regions, including the posterior and anterior insula, the superior temporal gyrus, the secondary somatosensory cortex, the superior temporal cortex, the hippocampus, and the cingulate cortex^20–28^. More recent studies employing functional and structural connectivity approaches have further demonstrated that these regions are densely interconnected^29,30,26^. However, due to the temporal limitations of fMRI and PET, the dynamic interactions between regions within this richly connected cortical network during self-motion perception remain poorly understood. For example, Brandt and colleagues proposed a reciprocal inhibition mechanism between vestibular and visual cortices, whereby artificial vestibular stimulation reduces relative BOLD signal in posterior visual areas, and vice versa^31,32^. Others proposed that visual attention networks modulate activity within the insular vestibular cortex^33^.

These correlational approaches, such as fMRI and PET, face several important limitations. First, participants are typically in a supine, head-fixed position, and the vestibular stimulations used are neither natural nor physiologically equivalent to real-world motion^34,35^. Second, vestibular inputs are often dissociated from the visual and somatosensory inputs that normally accompany natural whole-body movements. Finally, caloric, galvanic, and acoustic vestibular stimulations also deliver extravestibular inputs, such as tactile, thermal, nociceptive, or auditory stimuli^34,35^, which are difficult to disentangle despite efforts to better control for these confounding factors (e.g., skin anesthesia at the ear level^26^).

Together, these limitations underscore the need for approaches that move beyond correlation and directly test the causal contribution of specific brain regions to self-motion perception. Invasive electrical brain stimulation (EBS) is widely considered as the gold-standard approach for functional brain mapping, providing the highest level of causal evidence for the functional involvement of specific brain regions or functional networks^36–38^. Pioneering EBS studies during awake brain surgeries have shown that stimulation of the parietal cortex or the superior temporal gyrus can elicit conscious perceptions of self-motion, such as rotations, vertigo, or floating sensations^39,40^. More recent case series in patients with epilepsy indicated that such sensations can be evoked primarily through EBS of the perisylvian temporal cortex^41^, the insular cortex^42^, the posterior parietal cortex^43^, and the cingulate cortex^44,45^ (for reviews, see Ref.^46–48^). These findings provide causal confirmation of the regions identified in nonhuman primates and in human non-invasive neuroimaging studies.

Beyond mere mapping of stimulated regions, EBS offers a unique opportunity to investigate the functional networks involved in the construct of conscious self-motion perception. By assessing EBS-evoked changes directly from intracranial stereoelectroencephalography (SEEG), this approach allows to characterize the spatiotemporal dynamics of brain networks associated with the emergence of conscious self-motion experience, as previously done for other EBS-evoked sensations^49,50^. Recent evidence indicates that crosstalk and connectivity between functional brain networks^51^ is an important mechanism for sensorimotor processing and self-perception^45,52^, and alterations in these patterns of connectivity contribute to related clinical conditions^53–55^. Understanding these networks may also have clinical implications, particularly for conditions involving altered self-motion perception, such as vestibular epilepsy^56^ or functional neurological disorders^57^.

To this end, we conducted a retrospective analysis in a large cohort of patients (*n* = 354) who underwent presurgical evaluation for drug-resistant epilepsy. These patients were implanted with intracerebral SEEG electrodes and received EBS during SEEG recordings. We analyzed the self-motion sensations elicited by EBS outside lesioned regions and areas without epileptogenic activity^36,45^, along with concurrent SEEG signal recordings. To our knowledge, this is the first study to combine EBS with SEEG to map the causal and dynamic contributions of cortical networks to self-motion perception in humans. This approach allows us to extend our findings to the understanding of neurotypical brain function, thereby opening the door to a better comprehension of the functional organization of the brain networks involved in self-motion perception.

## Material and methods

### Patients

We retrospectively analyzed data from 354 patients (176 females; mean age ± standard deviation (SD): 25.5 ± 14 years) with drug-resistant focal epilepsy who underwent invasive SEEG at our tertiary epilepsy center between August 2000 and December 2021. All patients had previously undergone a comprehensive presurgical evaluation, including detailed medical history, neurological examination, neuropsychological evaluation, brain MRI, and surface EEG. SEEG was performed exclusively for clinical purposes to delineate the epileptogenic zone and its relationship with eloquent cortices, as part of the standard presurgical workflow^58^. EBS was conducted to trigger habitual seizures and for functional mapping, following established clinical protocols^59,60^.

### Ethical approval

This retrospective study was approved by our Institutional Review Board (APHM, #PADS24-22_dgr). All patients, or their parents/guardians, provided written informed consent for SEEG investigations and the use of their data for clinical and research purposes, in accordance with the Declaration of Helsinki.

### Electrodes implantation and SEEG recordings

Electrode implantation was performed solely for clinical evaluation, and no additional electrodes were implanted for research purposes. Implantation schemes were determined during a multidisciplinary meeting involving epileptologists and neurosurgeons to sample the hypothesized epileptogenic zone, identified through non-invasive investigations, while also considering anatomical and vascular constraints. SEEG exploration was performed using 9- to 15-contact depth electrodes (Dixi Medical or Alcis; contact length: 2 mm, diameter: 0.8 mm, inter-contact spacing: 1.5 mm). Among the 354 patients, 49 underwent left hemispheric implantations, 40 right hemispheric implantations, and 265 bilateral implantations. The number of implanted electrodes did not differ significantly between hemispheres (left: mean ± SD = 6 ± 5; right: mean ± SD = 6 ± 5; *t* = 0.96, *p* = 0.338).

Stereo-EEG signals were recorded using 128- or 256-channel Natus systems at sampling rates of 256, 512, or 1024 Hz and stored without digital filtering. During data acquisition, two hardware filters were used: a high-pass filter (cut-off frequency: 1 Hz at 3 dB) and an antialiasing low-pass filter (cut- off frequencies: 97 Hz for 256 Hz sampling, 170 Hz for 512 Hz sampling, and 340 Hz for 1024 Hz sampling). Two adjacent contacts on a SEEG electrode located in the white matter, distant from the presumed epileptogenic zone, were used as the reference electrode.

### Electrical brain stimulation procedures

EBS was conducted with patients in either sitting or supine positions, instructed to keep their eyes open and remain relaxed. Occasionally, patients were asked to perform tasks such as reading a text, repeating words, counting, or maintaining their arms in a stretched and lifted position, as is standard practice in functional brain mapping during EBS^38^. EBS was delivered using a current-regulated neurostimulator (Inomed, Colmar, France) in accordance with French SEEG guidelines^58^.

Square current pulses were administered between adjacent electrode contacts using two stimulation paradigms: (1) High-frequency stimulation at 50 Hz with a pulse width of 1ms and train duration of 3−8 s; or (2) Low-frequency stimulation at 1 Hz with a single pulse width of 2ms and train duration of 20−60s. These parameters were selected to minimize the risk of tissue damage, by maintaining the charge density per pulse below 50 μC/cm^2^ (Ref.^61^). Stimulation intensity was incrementally increased until either a sensation or an after-discharge was elicited, with a maximum intensity of 5 mA. However, intensities above 3mA were rarely applied. Each patient received a total of 1 to 175 high-frequency EBS (mean ± SD: 31 ± 25 EBS), and 1 to 185 low-frequency EBS (mean ± SD: 25 ± 33 EBS).

Importantly, patients were not informed about which contacts were being stimulated or when stimulation occurred. After each EBS, patients were systematically asked: “Did you feel anything?” to ensure unbiased reporting of subjective experiences. Patients were asked to provide detailed reports on any sensations or symptoms elicited by the stimulation. Both SEEG responses, including after-discharges, and clinical effects were meticulously documented in specialized notebooks and/or directly on SEEG recordings using markers and text annotations. Video and SEEG recordings of instances where responses were evoked were archived for further analysis.

### Retrospective analysis of EBS-evoked responses

We retrospectively reviewed clinical records of patients’ subjective reports (somatosensory, visual, auditory, or other sensations), behavioral responses, clinical signs (e.g., muscle contraction), and changes in SEEG activity elicited by EBS. In total, we identified 2,197 responses evoked by high-frequency EBS (intensity range: 0.2−5 mA) and 819 responses evoked by low-frequency EBS (intensity range: 0.2−5 mA). Data were compiled into a structured database (Microsoft Excel, version 16.79.2), including stimulation parameters (frequency, intensity) and the bipolar electrode contacts involved.

Inclusion criteria for this retrospective study were: (1) EBS-evoked vestibular self-motion illusions, defined as sensations of vertigo, dizziness, whole-body rotation or translation, in the absence of physical motion, consistent with previous functional mapping studies of vestibular sensations^41,42^. We distinguished between relatively nonspecific vestibular experiences (vertigo and dizziness) and more specific self-motion percepts (whole-body rotation and translation). Whereas vertigo and dizziness reflected a general disturbance of self-motion perception or spatial orientation (e.g., head spinning or feeling dizzy), rotation and translation involved explicit perceptions of angular or linear displacement of the body in space (e.g., floating, falling, spinning).; (2) EBS localized outside of both lesion or epileptogenic zones; (3) the absence of EBS-inducted habitual auras or seizures.

Responses were verified on video-SEEG recordings by a neurologist (S.L.) and two neuroscientists (Z.D. and C.L.) to ensure accuracy and consistency in reporting.

### Anatomical localization of EBS-evoking self-motion illusions

Electrode contacts where EBS elicited illusory self-motion sensations were localized with two distinct levels of spatial precision, depending on the period of implantation (**Figure 1**).

**Figure 1.**
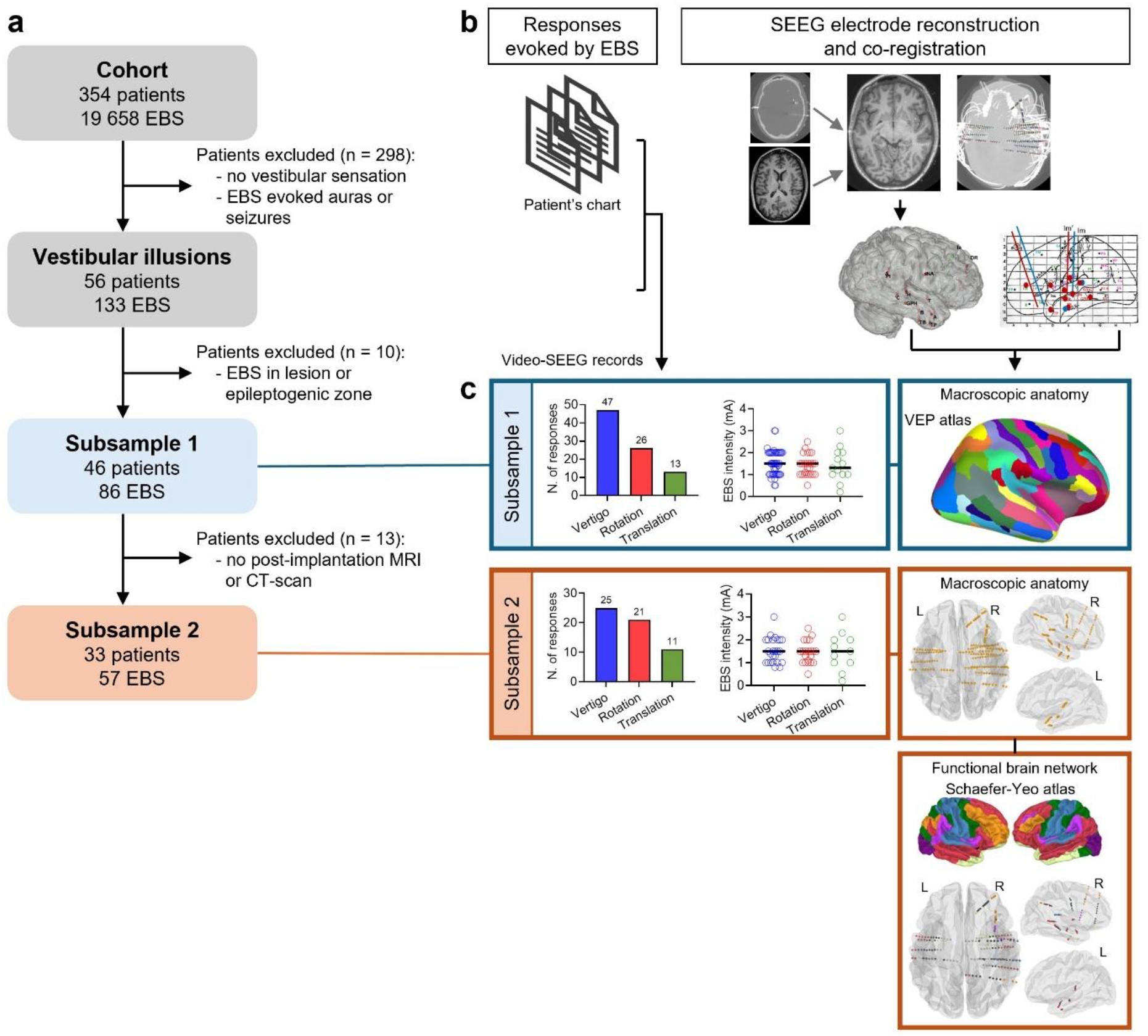
Procedures for the retrospective cohort analysis. **(a)** Flowchart of patient inclusion based on vestibular self-motion sensations evoked by electrical brain stimulation (EBS). **(b)** Data from medical records and results from video-stereo-electroencephalography (SEEG) investigations were retrospectively reviewed to identify EBS-evoked self-motion sensations, which were confirmed through visualization of video recordings. The localization of implanted brain electrodes was determined based on implantation schemes and macroscopic anatomical landmarks for patients implanted in earlier years (Subsample 1, *n* = 46) or precisely localized via co-registration of post-implantation CT scans and postoperative MRI for more recently implanted patients (Subsample 2, *n* = 33). **(c)** For each subsample, the number of self-motion responses evoked by EBS is presented, stratified by sensations of vertigo/dizziness, rotation, and translation, along with the stimulation intensity parameters (in mA) that triggered these responses.

For patients implanted after 2010, we performed automatic localization and labelling of each electrode contact using the open-source graphical user interface GARDEL (https://meg.univamu.fr/doku.php?id=epitools:gardel)^62^. This process involved co-registering pre-implantation MRI and post-implantation CT scans, followed by automatic recognition of each electrode contact and precise anatomical localization, resulting in patient-specific 3D maps of electrode contacts. Using GARDEL, each electrode contact was automatically assigned to its corresponding anatomical region by projecting the Virtual Epileptic Patient (VEP) atlas^63^ (which includes 162 brain regions based on parcellation from the Destrieux Atlas^64^, obtained after FreeSurfer segmentation^65^). The anatomical labelling of all electrode contacts was manually verified on the MRI by an expert neurologist (S.L.). For group-level analyses, electrode coordinates in the patient’s brain were converted to MNI space using custom scripts in Matlab 2020 (The MathWorks, Inc., USA) implemented in GARDEL and SPM 12^66^. The spatial distribution of electrodes implanted in the included patients is shown in **Figure 1a**.

For patients implanted before 2009, post-implantation CT and/or MRI scans were no longer available. In these cases, EBS localization was retrospectively extracted from SEEG clinical reports, in which electrode positions had been determined by clinicians based on the original implantation plans and anatomical targeting schemes. Anatomical labelling was performed according to the VEP atlas parcellation^63^ and used for quantitative group analyses only.

Following anatomical localization, cortical contacts were assigned to the corresponding 7 networks from the Schaefer-Yeo atlas^51,67^ following established procedures (https://github.com/ThomasYeoLab/CBIG/tree/master/stable_projects/brain_parcellation/Schaefer2018_LocalGlobal/Parcellations/project_to_individual). The 7-network Schaefer-Yeo atlas was projected from the *fsaverage* template onto each individual cortical surface using FreeSurfer (version 8.2.0). The atlas was mapped separately onto the left and right hemispheres using Freesurfer tools. The resulting patient-specific surface annotations were converted into volumetric parcellations in individual anatomical space. Each SEEG contact was then assigned to the Schaefer network corresponding to the nearest voxel to its coordinates using custom python script. Contacts located outside the cortical parcellation were classified as white matter, hippocampal, or subcortical contacts based on their anatomical localization (**Figure 2a**).

**Figure 2.**
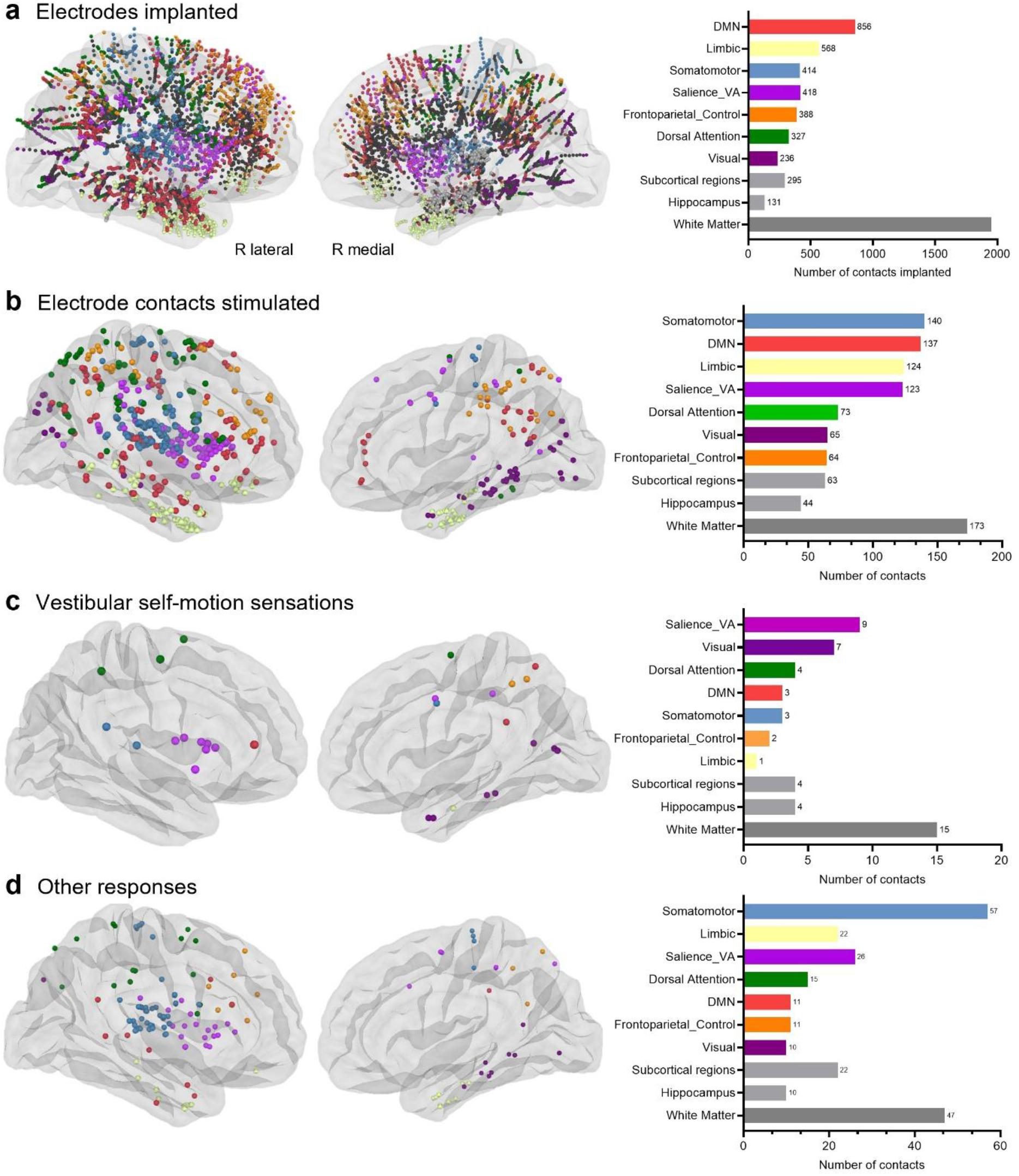
Spatial distribution of electrode contacts across functional networks from the Schaefer atlas in Subsample 2. **(a)** Spatial distribution of all implanted SEEG contacts across participants (*n* = 33 patients; 5,503 contacts). **(b)** Distribution of stimulation sites located outside the epileptogenic and lesioned zones (*n* = 181 cortical contacts assigned to functional networks). **(c)** Spatial distribution of stimulation sites eliciting vestibular sensations (n = 29 contacts assigned to functional networks). **(d)** Spatial distribution of stimulation sites eliciting non-vestibular responses (n = 152 contacts assigned to functional networks). Contacts are color-coded according to their assignment to the Schaefer sevennetwork atlas. Bar plots indicate the number of contacts within each functional network. Subcortical contacts, hippocampal contacts, and contacts located in white matter are displayed only in the bar plot. VA: ventral attention network; DMN: default mode network.

As stimulation was delivered in a bipolar configuration between pairs of adjacent contacts, the mean coordinate of the two contacts was used as the representative location for spatial analyses and visualization. For visualization purposes, left-hemisphere electrode reconstructions were flipped, mapping all electrodes to the right cerebral hemisphere. Cortical contacts were visualized on the MNI *fsaverage* template using the MNE toolbox (version 1.9.0)^68^.

## Results

### Self-motion sensations evoked by EBS

Vestibular self-motion sensations were evoked in 46 patients (21 females; mean age ± SD: 25 ± 11 years) during 86 EBS, all of which were applied outside epileptogenic and lesioned regions. Vestibular sensations were a relatively rare perceptual phenomenon, representing only 3.9% of all perceptual and behavioral responses elicited by EBS across the entire patient cohort. The distribution of evoked vestibular self-motion sensations and associated EBS intensity parameters is summarized in **Figure 1c**.

The majority of the elicited vestibular sensations were nonspecific self-motion experience, such as vertigo, dizziness, and instability (*n* = 47, 54.7%). In contrast, more specific self-motion experience were reported following 39 EBS. Rotational sensations were elicited by 26 EBS (30.2%), primarily in the yaw plane (*n* = 11), and less frequently in the pitch (*n* = 5) and roll (*n* = 3) planes. Notably, rotational sensations were predominantly in the clockwise direction (*n* = 11), compared to counterclockwise (*n* = 1) and swaying (*n* = 4). Sensations of body translation were less frequently elicited (*n* = 13, 15.1%), with upward motion and floating sensations (*n* = 8) reported more often than sensations of falling (*n* = 2).

These vestibular sensations were predominantly elicited by high-frequency EBS at 50 Hz (*n* = 72, 83.7%), whereas only a minority were evoked by low-frequency EBS at 1−2 Hz (*n* = 14, 16.3%). Analysis of stimulation intensity revealed that different vestibular sensations were not associated with differences in EBS intensity (*F*_(2,83)_ = 0.462, *p* = 0.632; vertigo/dizziness (mean ± SD): 1.5 ± 0.6 mA; rotations: 1.4 ± 0.5 mA; translations: 1.4 ± 0.8 mA). Furthermore, the thresholds for eliciting vestibular sensations (1.5 ± 0.5 mA; range: 0.2–3 mA) did not differ significantly from the thresholds for eliciting other perceptual and behavioral responses in the same 46 patients (1.7 ± 0.6 mA; range: 0.2–3 mA; *t*_(284)_= 1.196, *p* = 0.233).

In most cases (*n* = 51, 59.3%), vestibular sensations were not accompanied by other responses. However, in addition to vestibular sensations, some patients reported unpleasant sensations, fear, or anxiety (*n* = 7), neurovegetative responses (e.g., nausea, palpitations, rising epigastric sensation, lacrimation) (*n* = 7), somatosensory or body image distortions (*n* = 6) or exhibited motor or oculomotor responses (*n* = 5). Other accompanying sensations included perceived change in body temperature (*n* = 4), visual sensation (*n* = 4), derealization (*n* = 4), headache (*n* = 2), auditory sensation (*n* = 1), and dysarthria (*n* = 1).

### Localization of electrode contacts eliciting self-motion sensations

A global analysis of vestibular self-motion sensations in the largest patient sample (Subsample 1: *n* = 46, 86 EBS), combining implantation scheme-based localization for earlier patients (*n* = 13) with MRI/CT-based electrode localization for more recent patients (*n* = 33), revealed a wide anatomical distribution of electrode contacts. Responses were equally distributed across hemispheres (43 left, 43 right EBS) and originated predominantly from frontal (29.1%), temporal (24.4%), parietal (20.9%), and insular (15.1%) cortices, with fewer responses originating from occipital (4.7%) and subcortical (5.8%) structures.

A detailed analysis of electrode contact coordinates was performed in Subsample 2 (*n* = 33, 57 EBS), in which all electrodes were precisely localized through co-registration with post-implantation MRI or CT scans. In total, 5,503 contacts were implanted, including 2,462 in the left hemisphere and 3,041 in the right hemisphere. Of the 1,006 contacts stimulated outside the epileptogenic and lesioned zones, 283 EBS evoked responses, whereas 723 did not. Among these response-evoking EBS, 57 elicited vestibular self-motion sensations. Four patients experienced vestibular sensations during repeated stimulation of the same site, yielding a total of 52 unique EBS sites associated with vestibular self-motion sensations.

Vestibular self-motion sensations were broadly distributed across cortical and subcortical regions but showed preferential clustering within the posteromedial cortex, insula, and medial temporal lobe. When normalized by the total number of EBS delivered within each region, the highest vestibular elicitation rate was observed in the posteromedial cortex (18.4%, 7/38 EBS), comprising the precuneus and posterior cingulate cortex. Elevated response rates were also observed within the insula (7.1%, 8/112 EBS), particularly within its anterior part (4.5%, 5/112), and in medial temporal lobe, including the hippocampus (8.8%, 3/34 EBS) and rhinal cortex (8.8%, 3/37 EBS). Additional vestibular self-motion sensations were evoked by EBS in the middle cingulate, the amygdala, the inferior frontal gyrus, and premotor cortices.

Sensations of dizziness/vertigo exhibited the broadest anatomical distribution, including the hippocampal-parahippocampal complex, posterior insula, sensorimotor cortex (precentral and postcentral gyri), and inferior frontal gyrus. By contrast, sensations of rotations were preferentially associated with stimulation of posteromedial regions, particularly the precuneus and posterior cingulate cortex, as well as in the anterior insula. Translation sensations showed the most spatially restricted pattern, converging within the hippocampal-parahippocampal complex and anterior insula.

### Relation to functional brain networks

Mapping vestibular responses and other EBS-evoked responses onto the 7 functional networks from the Yeo-Schaefer atlas^67^ revealed a non-uniform distribution of responses across networks (*χ*^2^_(6)_ = 65.32, *p* < 0.0001). Among the 29 vestibular sites that fell into parcels from the atlas (**Figure 2c**), the largest proportion was found in the ventral attention network (31.0%, 9/29 EBS), followed by the visual (24.1%, 7/29 EBS) and dorsal attention (13.8%, 4/29 EBS) networks (**Figure 3a, Table 1**).

**Table 1.** Number and elicitation rate of vestibular sensations across 7-Schaefer networks. Vestibular rates represent the proportion of stimulation within each network that elicited a vestibular sensation. Confidence intervals (95% CI) were estimated using bootstrap resampling (10,000 iterations).

|  |  | Salience /<br>Ventral<br>attention | Visual | Dorsal<br>attention | Somatomotor | Default mode | Frontoparietal | Limbic | Total |
| --- | --- | --- | --- | --- | --- | --- | --- | --- | --- |
| EBS<br>delivered | *n* | 123 | 65 | 73 | 140 | 137 | 64 | 124 | 726 |
| Evoked<br>responses | Vestibular<br>*n*<br>% | 9<br>31.0% | 7<br>24.1% | 4<br>13.8% | 3<br>10.3% | 3<br>10.3% | 2<br>6.9% | 1<br>3.4% | 29<br>100% |
|  | Other<br>*n*<br>% | 26<br>17.1% | 10<br>6.6% | 15<br>9.9% | 57<br>37.5% | 11<br>7.2% | 11<br>7.2% | 22<br>14.5% | 152<br>100% |
|  | Total<br>*n*<br>% | 35<br>19.3% | 17<br>9.4% | 19<br>10.5% | 60<br>33.1% | 14<br>7.7% | 13<br>7.2% | 23<br>12.7% | 181<br>100% |
| Elicitation<br>rate per<br>network | Vestibular<br>%<br>95% CI | 7.3% *<br>[3.252, 12.195] | 10.8% *<br>[4.615, 18.462] | 5.5% *<br>[0.000, 0.050] | 2.1%<br>[0.000, 5.000] | 2.2<br>[0.000, 5.109] | 3.1%<br>[0.000, 7.813] | 0.8%<br>[0.000, 2.419] |  |
|  | Other<br>%<br>95% CI | 21.1% *<br>[13.821, 28.455] | 15.4% *<br>[7.692, 24.615] | 20.5% *<br>[12.329, 30.137] | 40.7% *<br>[32.857, 48.571] | 8.0% *<br>[3.650, 13.139] | 17.2% *<br>[7.813, 26.563] | 17.7% *<br>[11.290, 25.000] |  |

**Figure 3.**
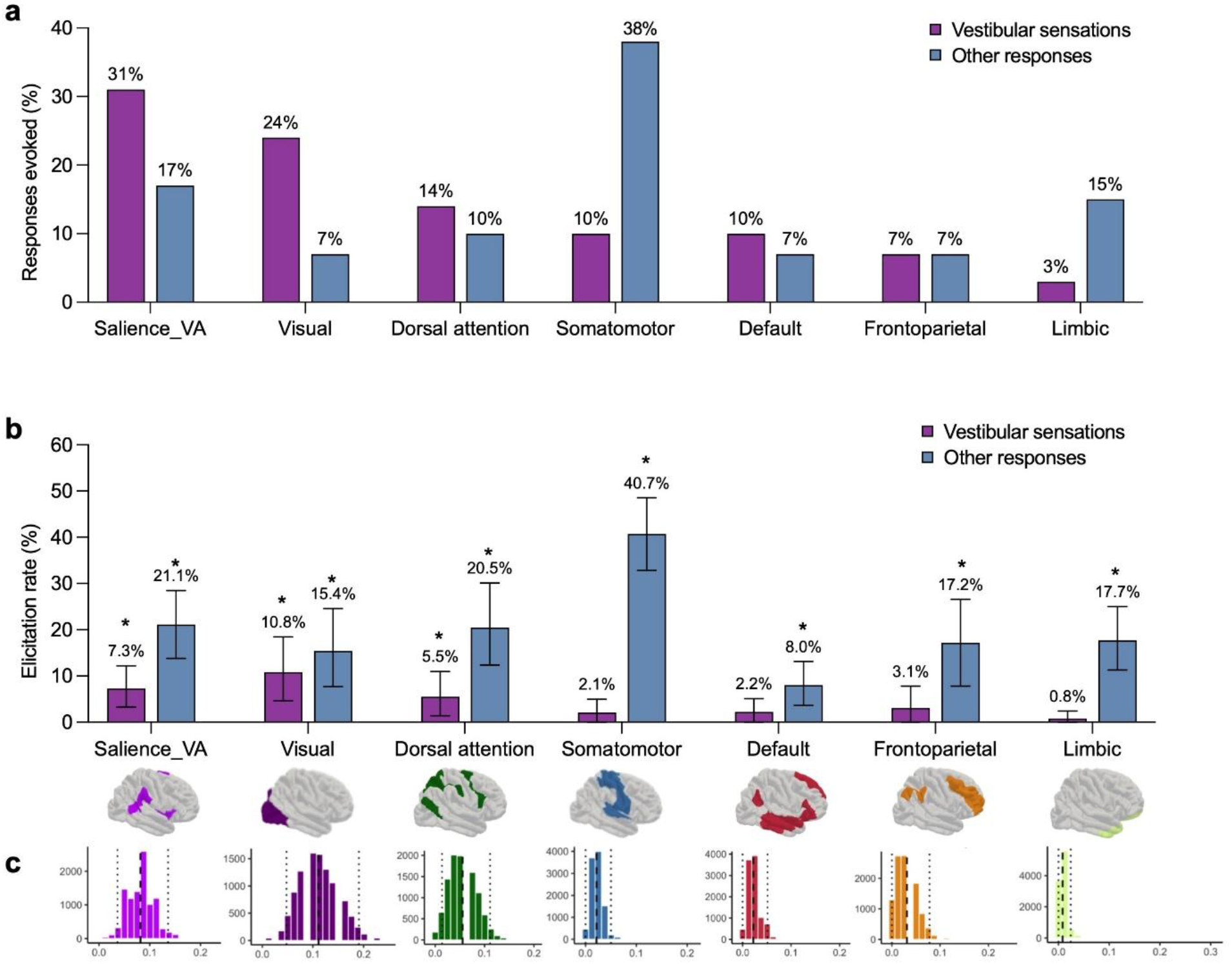
Distribution of EBS-evoked responses across functional networks from the Schaefer atlas. **(a)** Distribution of vestibular self-motion sensations (*n* = 29) and other EBS-evoked responses (*n* = 152) across the seven functional networks, ordered by proportion of vestibular sensations within each network. **(b)** Elicitation rate (number of responses / number of EBS per functional network) of vestibular sensations and other EBS-evoked responses in each functional network. Error bars represent 95% confidence intervals (CI) estimated from bootstrap distributions (10,000 iterations). **(c)** Observed elicitation rates (dashed black line) within 95% bootstrap CIs (dotted lines) for vestibular sensations. VA: ventral attention network.

To determine whether vestibular sensations were preferentially associated with specific functional networks, vestibular sensation labels were randomly reassigned across all stimulated sites while preserving both the total number of vestibular (*n* = 29) and other (*n* = 152) responses. A permutation analysis (10,000 iterations) showed that the observed distribution of vestibular sensations across functional networks differed significantly from that expected based on the sampling density of stimulated sites (*χ*^2^_(6)_ = 15.83, *p* = 0.005). Vestibular sensations occurred more frequently than expected by chance within the visual network (7 observed vs. 2.72 expected responses; residual = 2.59), the ventral attention network (9 observed vs. 5.61 expected responses; residual = 1.43), the dorsal attention network (4 observed vs. 3.04 expected responses; residual = 0.55) and the default mode network (3 observed vs. 2.24 expected responses; residual = 0.51).

Bootstrap-derived elicitation rates, which account for the number of EBS per functional network, revealed significantly higher elicitation rates specifically in the visual network (10.8%, 95% CI [4.615, 18.462]) and ventral attention network (7.3%, 95% CI [3.252, 12.195]), followed by the dorsal attention network (5.5%, 95% CI [1.370, 10.959]) (**Figure 3b, Table 1**). In contrast, elicitation rates for other responses were significant in all 7 functional networks (**Table 1**).

## Discussion

In this study, we report EBS findings from 86 stimulations obtained in 46 patients with drug-resistant focal epilepsy who experienced self-motion sensations, including rotation, translation, dizziness and vertigo. The diverse anatomical distribution of these responses supports the view that vestibular self-motion perception emerges from activity within a wide cortical network^4,5^. Importantly, the network distribution differed between vestibular and other types of EBS responses. Responses evoking other types of sensations (motor, language, somatosensory, visual, etc.) were predominantly distributed across the somatomotor, salience/ventral attention, limbic, and dorsal attention networks, with comparatively fewer responses arising from the default mode, frontoparietal, and visual networks. This pattern is consistent with a previous large-scale EBS analysis^69^, in which the highest elicitation rates were observed in the somatomotor and visual networks, followed by the salience and dorsal attention networks. In contrast, we found that vestibular self-motion sensations showed a more specific networklevel distribution, with the highest elicitation rates observed in the salience/ventral attention, visual, and dorsal attention networks, and comparatively fewer responses arising from the somatomotor, frontoparietal, and limbic networks. These findings suggest that the vestibular self-motion network preferentially involves cortical networks involved in attentional control and multisensory processing. We discuss the contribution of these functional networks below.

### Contributions of the salience/ventral attention network to self-motion perception

One of the main findings of the present study was the preferential involvement of the salience/ ventral attention network in vestibular self-motion sensations. Under normal conditions, vestibular signals contribute to the pre-reflective consciousness of the body and its movement/orientation in space, forming an implicit component of the minimal self^70^. EBS may disrupt this normally seamless integration, transforming an implicit bodily signal into a salient and conscious experience of self-motion. In this context, the salience network, comprising key hubs in the anterior insula and anterior cingulate cortex and extending to subcortical structures such as the amygdala and thalamus, may contribute to bringing altered bodily signals into conscious perception. This idea is supported by the proposed role of the salience network in detecting behaviorally relevant or unexpected changes in internal and external states and allocating attention toward them^71,72^. This interpretation converges with the recently described action-mode network (AMN), anatomically close to the salience/ventral attention network^72,73^, which includes an AMN-Bodily Self subnetwork^73^. This subnetwork may provide an interface between the detection of salient bodily changes and their integration into the subjective experience of the bodily self.

Several vestibular self-motion sensations were evoked by EBS of the insula. As a key hub of the salience/ventral attention network^74^ and a core area of the cortical vestibular system^5,18,25^, the insula may represent an important entry point through which EBS engages broader networks involved in the detection and integration of salient bodily changes. Although previous studies have emphasized a predominant contribution of the posterior insula^18,42,75^, vestibular self-motion sensations in our study were more frequently elicited by stimulation of anterior than posterior insular regions. This may be relevant to the subjective dimension of the elicited sensations, as the anterior insula has been proposed to integrate bodily and interoceptive signals with affective and attentional processes^76,77^ and to contribute to the representation of ongoing bodily states and movements^77^. This might be related to unpleasant feelings, fear, and nausea that were in some instances associated with EBS in the anterior in the insula in our sample, as well as in previous case series^78^.

### Contributions of the dorsal attention network to self-motion perception

The preferential involvement of the dorsal attention network provides a complementary perspective on the neural mechanisms underlying vestibular self-motion perception. This network, which includes the intraparietal sulcus, superior parietal lobule, and frontal eye fields^79^, is classically involved in the top-down orientation of attention toward behaviorally relevant sensory information and in the selection and maintenance of task-relevant representations^79–81^. The recruitment of this network in our study may therefore reflect the role of vestibular signals in informing the brain about changes in body orientation and movement, while engaging mechanisms involved in spatially directed attentional selection and orientation. We propose that the dorsal attention network may contribute to allocating attention toward the spatial implications of altered vestibular signals, in concert with the salience/ventral attention network, which detects salient changes in bodily state and facilitates their conscious perception. Once attention has been reoriented, the dorsal attention network could contribute to selecting and maintaining attention toward the spatial consequences of that altered signal. Consistent with this interpretation, experimental deprivation of vestibular otolithic information during parabolic flight has been shown to alter visuospatial attention, increasing automatic attention while reducing the voluntary maintenance of attention at a cued location^82^. Together, these findings suggest that vestibular self-motion processing is linked to attentional mechanisms.

### Contributions of the visual network to self-motion perception

EBS within the visual network elicited vestibular sensations, particularly dizziness and vertigo, suggesting that perturbation of the visual network can alter the perception of self-motion. Responses were predominantly associated with medial occipitotemporal regions, including the lingual and fusiform gyri, rhinal cortex, and collateral sulcus. These regions have been implicated in scene processing, spatial representations, and navigation. Along this line, visual cortical regions contribute directly to the processing of self-motion by integrating visual and vestibular information. Previous works in non-human primates showed that visual motion areas contain populations of neurons responsive to both optic flow and vestibular stimulation, contributing to multisensory self-motion processing^12,83^. Stimulation of these sites may alter neuronal processing in populations that integrate visual and vestibular signals and thereby alter the multisensory construct of self-motion.

Converging evidence in humans further supports this interpretation. Galvanic vestibular stimulation has been shown to increase functional connectivity within visual areas^84^, while patients with vestibular agnosia, characterized by impaired conscious perception of self-motion, showed altered functional connectivity in the visual network regions compared to patients with preserved vestibular self-motion perception^85^. Finally, causal evidence comes from EBS in visual and temporo-occipital regions, including the middle temporal and lingual gyri, which can evoke sensations such as rotation, dizziness, or levitation^41^.

The contribution of visual regions may also be understood as reflecting reciprocal inhibitory mechanisms between visual and vestibular cortical systems. When visual processing is prioritized, BOLD signal within the PIVC is inhibited, potentially reducing the influence of competing vestibular signals during visual processing^31,86^. This interaction appears to be bidirectional, as vestibular stimulation has also been shown to suppress BOLD signal within visual areas^32^. Thus, EBS in sites within the visual network may disrupt this balance and alter multisensory processing, potentially leading to erroneous self-motion perception.

### Contributions of the hippocampus to self-motion perception

Although the hippocampus was not included in the functional network parcellation we used^51^, we note that several EBS in the hippocampus evoked vestibular self-motion sensations. Some case series have also reported sensations of whole-body translations and vertigo/dizziness during EBS of the hippocampus^41,87^. The hippocampus is not typically associated with vertigo and dizziness, but it is often linked to spatial disorientation and spatial cognition deficits^88^. Single-cell recordings in the hippocampus show that this structure encodes real-world whole-body displacements^13^ or virtual reality navigation^89^. Furthermore, hippocampal volume has been shown to decrease in cases of peripheral vestibular dysfunction^90^. Interestingly, the hippocampus is strongly connected to core areas of the cortical vestibular network, such as the insula and secondary visual areas^91–93^, and EBS in the hippocampus may evoke self-disorientation.

### Limitations of the study

Several limitations should be considered when interpreting our findings. First, the anatomical distribution of EBS sites was determined by clinical needs, resulting in uneven sampling across cortical regions and potential sampling bias. Although our analyses accounted for the number of stimulations delivered within each region and functional network, uneven anatomical coverage remains an inherent limitation of intracranial EBS studies. Second, our network-level analysis was restricted to cortical regions represented in the Schaefer atlas^67^ and therefore did not include subcortical structures or the hippocampus. This is particularly relevant given that several vestibular responses were elicited from medial temporal regions, including the hippocampus. Third, EBS can recruit connected neuronal populations and propagate through anatomically and functionally connected pathways. Consequently, vestibular sensations elicited by stimulation of a given region may reflect perturbation of a broader network rather than the function of the stimulated site alone. Future studies combining EBS with analysis of functional connectivity could help address this issue by characterizing how stimulation-induced vestibular illusions alter interactions between cortical networks. Finally, the retrospective nature of the study and the clinical context limited the systematic characterization of subjective experiences. Vestibular sensations were based on spontaneous patient reports. Consequently, the phenomenological characterization of potentially complex self-motion experiences was potentially limited. Prospective studies using standardized assessments could provide a more detailed characterization of different selfmotion perceptions.

## Conclusions

Results from the present study demonstrate that self-motion perception arises from the dynamic interaction of distributed functional brain networks, particularly the salience/ventral attention, dorsal attention, and visual networks, rather than isolated vestibular regions. These results provide causal evidence that perturbations in these networks can generate illusory self-motion sensations, offering new insights into the neural mechanisms underlying vestibular epilepsy, functional neurological disorders characterized by dizziness, and spatial disorientation.

## Acknowledgements

This work was supported by the ANR VESTISELF project, grant ANR-19-CE37-0027 of the French Agence Nationale de la Recherche to C. Lopez and F. Bartolomei.

## Use of AI

During the preparation of this manuscript, the authors used Vibe (Mistral Medium 3.5, Mistral AI) to improve the readability, clarity, and language of the text.

